# A meta-analysis of neural systems underlying delay discounting: Implications for transdiagnostic research

**DOI:** 10.1101/2022.10.12.511959

**Authors:** Nikhil V. Lakhani, Min K. Souther, Bema Boateng, Joseph W. Kable

## Abstract

Delay discounting is a promising paradigm for transdiagnostic research because both excessive and insufficient tendency to discount future rewards have been reported across diagnoses. Because delay discounting involves multiple neurocognitive functions, researchers have used many strategies to characterize brain activity during delay discounting. However, which of these analytic approaches yield truly robust and replicable findings remains unclear. To this end, we conducted a meta-analysis of 80 fMRI studies of delay discounting, testing which statistical contrasts give rise to reliable effects across studies. Despite being a widely used analytic approach, comparing impulsive and patient choices did not reliably yield the expected effects. Instead, subjective value contrasts reliably engaged the valuation network, and task versus baseline and choice difficulty contrasts reliably engaged regions in the frontoparietal and salience networks. We strongly recommend that future neuroimaging studies of delay discounting use these analytic approaches shown to reliably identify specific networks. In addition, we provide all cluster maps from our meta-analysis for use as a priori regions of interest for future experiments.

## 1. INTRODUCTION

In 2009, the National Institute of Mental Health launched the Research Domain Criteria (RDoC) initiative to identify common biological mechanisms for understanding mental disorders (Insel, 2009). After over a decade of research, reliability remains a central challenge for identifying biobehavioral measures that provide novel insights into the core neurocognitive features exhibited across disorders (MacNamara & Phan, 2016). Delay discounting, or the tendency to devalue rewards that are distant in the future, is a promising candidate paradigm for identifying shared reward-related changes across diagnostic populations (Lempert et al., 2019). Currently categorized as a subconstruct of reward valuation under positive valence systems, delay discounting has been implicated in a wide range of psychiatric disorders, as both excessive discounting (e.g., substance use disorder, depression, schizophrenia, binge eating disorder) and insufficient discounting (e.g., anorexia) can be maladaptive (Amlung et al., 2017, 2019). In addition to this dimensional feature that cuts across diagnoses, delay discounting has been shown to be a strong prognostic indicator of treatment response (Washio et al., 2011) and relapse (Sheffer et al., 2014), and can be modified using pharmacological (Demoto et al., 2012; de Wit & Mitchell, 2010; Orsini et al., 2017), neural (Hecht et al., 2013), and behavioral (Ciaramelli et al., 2019; Lempert et al., 2017) interventions, which demonstrates its potential for clinical utility. Coupled with its high test-retest reliability (Enkavi et al., 2019) and evidence for a genetic basis (Anokhin et al., 2014; Gray et al., 2019), delay discounting indeed has many appealing features as a transdiagnostic construct.

Supplementing this emerging evidence for the validity of delay discounting as a transdiagnostic behavioral construct, a plethora of imaging studies have investigated the neural activity associated with delay discounting. Subsequently, early meta-analyses revealed reliable activation of brain regions linked to reward valuation and task engagement during delay discounting, including large swaths of medial and lateral prefrontal cortex, parietal cortex, posterior cingulate cortex (PCC), insula, and striatum (Carter et al., 2010; Wesley & Bickel, 2014). Furthermore, transdiagnostic meta-analysis suggested that brain activity during delay discounting varies by psychiatric diagnoses and symptoms (Chen et al., 2021). Notably, however, while researchers have used a wide variety of analytic strategies (i.e., statistical contrasts) to characterize brain activity during delay discounting, these early meta-analyses did not distinguish between these different strategies. It remains unclear which of these approaches yield truly robust and replicable findings. Recent research has shown that analytic choices can have a profound impact on the validity and interpretation of neuroimaging results (Botvinik-Nezer et al., 2020; Mumford et al., 2025). Identifying the neural effects that are reliable and reproducible across delay discounting experiments is an important foundation for future transdiagnostic neuroimaging research. Here, we used meta-analysis to systematically evaluate the reliability of existing analytic approaches.

One commonly used analytic approach is to characterize neural activity based on choice behavior. This is often accomplished by comparing brain activity when one chooses the smaller sooner reward (*SSR*) versus the larger later reward (*LLR*). This approach is motivated by a prominent hypothesis for neural mechanisms underlying maladaptive decision-making called the Competing Neurobehavioral Decision Systems (CNDS) model (Bickel et al., 2019). The CNDS hypothesis draws upon one of the first neuroimaging studies of delay discounting, which proposed two separate neural systems in the healthy brain: one system, composed of the striatum, ventromedial prefrontal cortex (vmPFC), and PCC, biases decisions toward the *SSR*; whereas another competing system, composed of the frontoparietal and salience networks, biases decisions toward the *LLR* (McClure et al., 2004). The CNDS hypothesis posits that impulsive choices are driven by bottom-up reward signals in response to immediate rewards, whereas patient choices are driven by top-down inhibitory control signals that can override this sensitivity. Taken together, both excessive and insufficient discounting can arise when competition between the bottom-up reward signals and top-down inhibitory signals is not balanced. This framework has become instrumental for subsequent clinical neuroscience research, as numerous studies have contrasted *SSR* and *LLR* choices to try and identify the neural systems underlying impulsivity and self-regulation features of various psychiatric disorders (Carlisi et al., 2016, 2017; Claus et al., 2011; Lim et al., 2017; Murphy et al., 2017; Norman et al., 2017; Wang et al., 2017).

Another commonly used analytic approach in the delay discounting literature is to characterize brain activity using latent constructs that guide choice behavior, such as subjective value and choice difficulty. In practice, this approach identifies brain activity that is parametrically correlated with a continuous variable that is either presented (e.g., dollar amount of reward, number of days to wait) or estimated based on choice behavior (e.g., subjective value, choice difficulty). This approach is grounded in another early neuroimaging study of delay discounting, which proposed that the striatum, vmPFC, and PCC, formed a unitary valuation system for both immediate and delayed rewards (Kable & Glimcher, 2007). In this view, rather than driving specific choice behavior, frontoparietal and salience regions are potentially recruited for domain-general task-related demands (Duncan & Owen, 2000; Duncan, 2010). This account is also consistent with previous meta-analytic work that found that the ventral striatum, vmPFC, and PCC encode positive effects of subjective value across multiple reward domains (Bartra et al., 2013; Clithero & Rangel, 2014). On this account, disruptions in value signals in either direction can be maladaptive, and reduced value response has been reported across disorders characterized by anhedonia or reduced motivation (Brown et al., 2020; Souther et al., 2022), whereas pathological gambling has been linked to exaggerated value signals (Miedl et al., 2012). However, although contrasts targeting value-related brain activity have been widely used in decision neuroscience, relatively fewer clinical neuroimaging studies of delay discounting have taken this approach (Miedl et al., 2012; Souther et al., 2022; Wiehler et al., 2017).

It is important to note that both analytic approaches converge on which brain regions are involved during delay discounting: the valuation (e.g., striatum, vmPFC, PCC), frontoparietal, and salience networks. However, what differentiates these approaches is how to best identify brain activity in these regions. The former uses binary comparisons between impulsive and patient choices, whereas the latter uses correlations with continuous attributes of choice options, such as subjective value, amount, delay, and choice difficulty.

In this study, we performed a series of meta-analyses of fMRI findings to identify which analytic approaches during delay discounting paradigms yield reliable findings across studies in the face of variability in study designs and the presence of psychiatric diagnoses. For each of these analytic approaches, we conducted a multilevel kernel density analysis (MKDA; Wager et al., 2007) to quantify the proportion of studies that report consistent effects near a given anatomical location. Our primary aim is to assess the reliability of the two most widely used analytic approaches, contrasting impulsive and patient choices and correlating activity with objective (magnitude, delay) or latent model-derived (subjective value) variables. For completeness, we also evaluate the reliability of other commonly reported contrasts, including task versus control, hard versus easy choices, and choices involving an immediate reward versus those between only delayed rewards.

To our knowledge, there is one published meta-analysis on delay discounting fMRI that aimed to systematically compare these contrasts (Schüller et al., 2019). However, the sample sizes in that paper ranged from 3 to 13 experiments, which does not meet the currently recommended guidelines for meta-analytic sample sizes (Eickhoff et al., 2016). Our meta-analyses include significantly more studies, yielding robust sample sizes (9 to 29 experiments per contrast, including >20 for the most critical contrasts) and permitting the examination of more contrasts (nine versus six). Importantly, with these methodological improvements, we come to a significantly different conclusion. Schüller et al. (2019) found that all analytic contrasts yielded significant clusters, suggesting equal adequacy of both the choice-based and latent construct-based approaches. On the contrary, with more than three times the sample size, we conclude that the expected effects of choice-based contrasts (both *SSR* and *LLR*) are not reliable. Given that choice-based contrasts are one of the most commonly used analytic approaches in existing studies, especially in experiments that focus on clinical populations, it is our hope that these more robust findings will help inform analytic decisions for future studies.

## 2. METHODS

Preregistration of the present meta-analysis is available on PROSPERO under protocol CRD42020150691.

### 2.1. Search strategy

We conducted a systematic literature search on PubMed and PsycInfo (final search on March 2, 2023). Based on previous meta-analyses of delay discounting (Amlung et al., 2019; Wesley & Bickel, 2014), we used the following search terms, which yielded 392 unique results: [(“Delay Discounting” OR “Temporal Discounting” OR “Delay of Gratification” OR “Intertemporal Choice” OR “Time Discounting”) AND (“fMRI” OR “Functional Magnetic Resonance Imaging”)]. We also reviewed the reference lists of previously published review articles and the results of searches for frequently occurring authors (i.e., 10 or more unique articles in the initial search results), which identified 2 additional articles.

### 2.2. Study selection

All articles identified through the above search strategy were evaluated according to the preregistered inclusion/exclusion criteria. For inclusion, the study had to meet the following criteria: (1) English or an English translation is available; (2) human participants with no brain lesions; (3) fMRI data collected while the participants were completing a delay discounting task in the scanner; (4) the task included a delay of at least 1 day and used monetary rewards, regardless of payment contingency; (5) conducted a parametric or categorical contrast analysis across the whole brain; (6) spatial coordinates are available in Talairach or Montreal Neurological Institute (MNI) space, either reported in the manuscript or provided by the corresponding author upon request; and (7) diagnosis of psychiatric disorders can be present.

Although existing meta-analyses of self-regulatory decision-making tend to focus on healthy adults or group comparisons (Han et al., 2018; Schüller et al., 2019; Yuan & Raz, 2014), inclusion of individuals with psychiatric disorders achieves two goals. First, it allows greater variability of the clinical or cognitive constructs (e.g., impulsivity, self-regulation, valuation, conflict monitoring, and action planning) that may be related to delay discounting. Second, given our ultimate aim is to identify specific neural systems engaged during delay discounting that may be related to transdiagnostic neurobehavioral processes, inclusion of psychopathology without regard to specific diagnoses seemed critical.

We excluded studies that (1) were not a primary source; (2) used non-monetary rewards; (3) had pharmacological manipulation unless they report effects in the placebo group; (4) were resting-state or volumetric analyses; (5) were connectivity or multi-voxel pattern analyses; (6) had overlapping samples with another included study. In the event of overlapping samples, we included the largest sample. To ensure inclusion criteria were standard and to limit bias, all studies were reviewed using Rayyan (https://rayyan.qcri.org; Ouzzani et al., 2016) by at least two independent reviewers (M.K.S. and J.W.K., M.K.S. and B.B., or all three). The inter-rater reliability was high (κ = 0.86; 93.9% agreement) and all disagreements were discussed and resolved by a consensus decision. A total of 80 studies (1311 foci; 3235 participants) were included in the quantitative meta-analysis (see Figure 1 for the PRISMA flow diagram; Table 1 for a list of studies and contrasts).

**Fig. 1.**
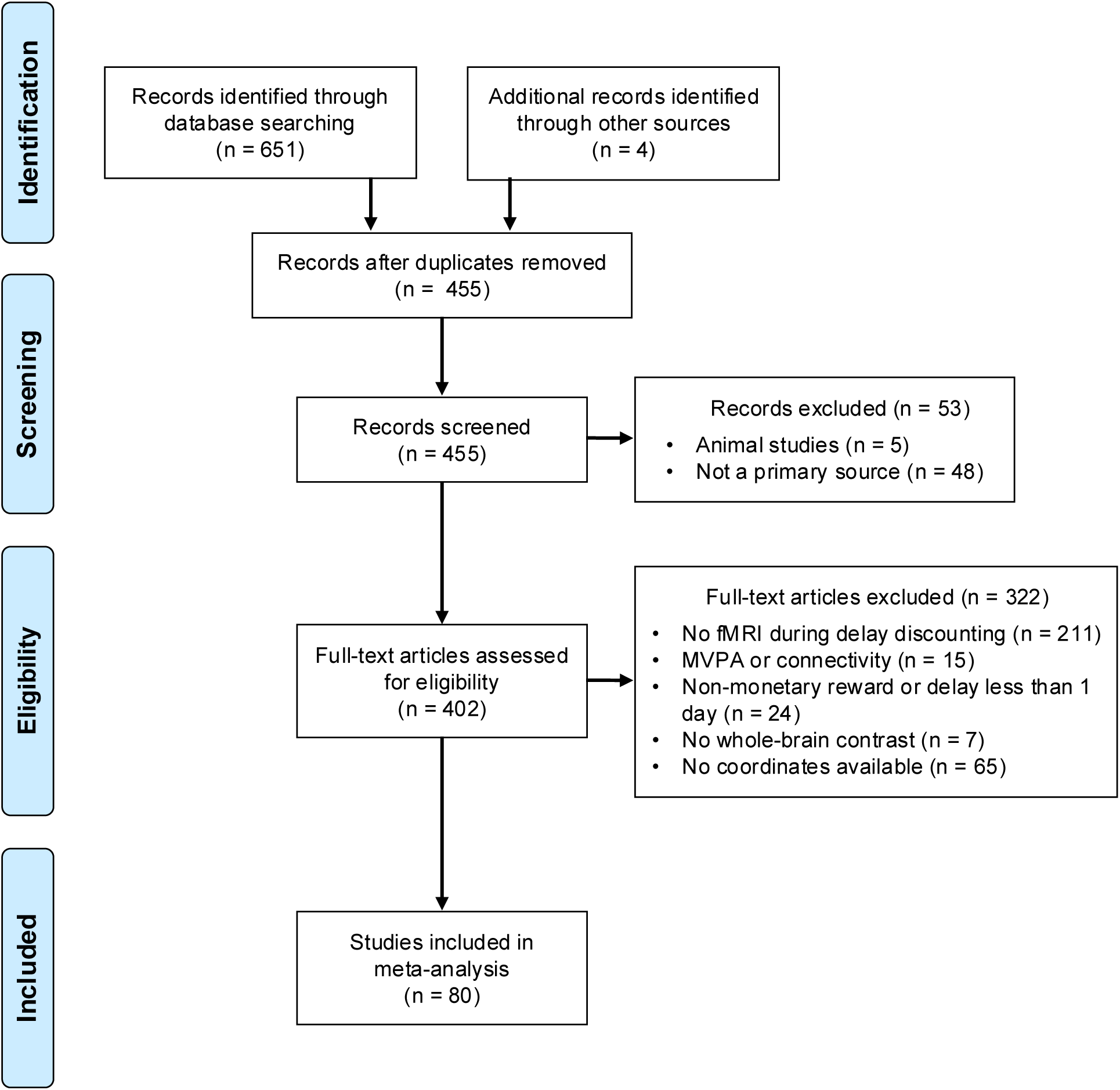
PRISMA flow diagram of study inclusion

**Table 1.**
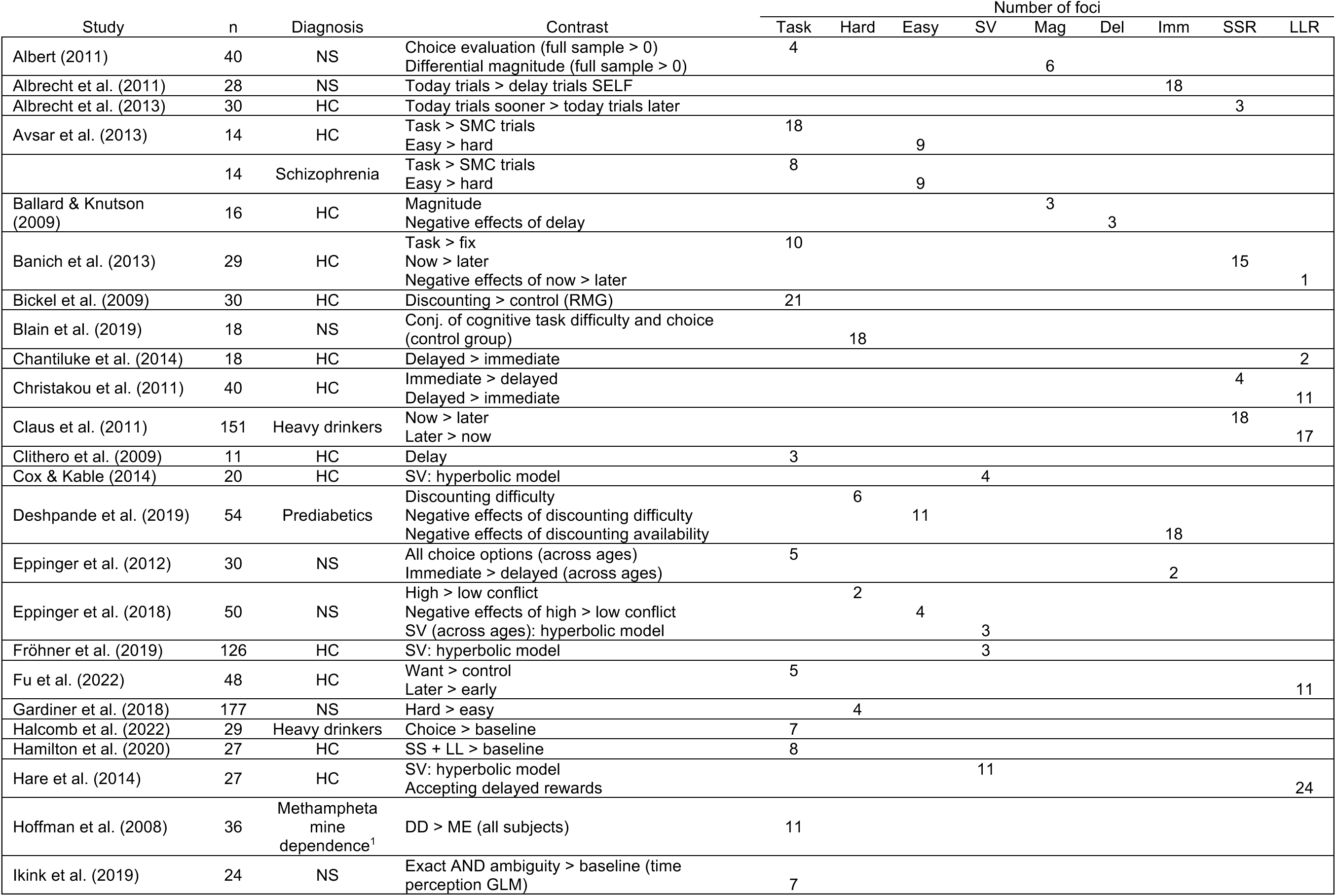
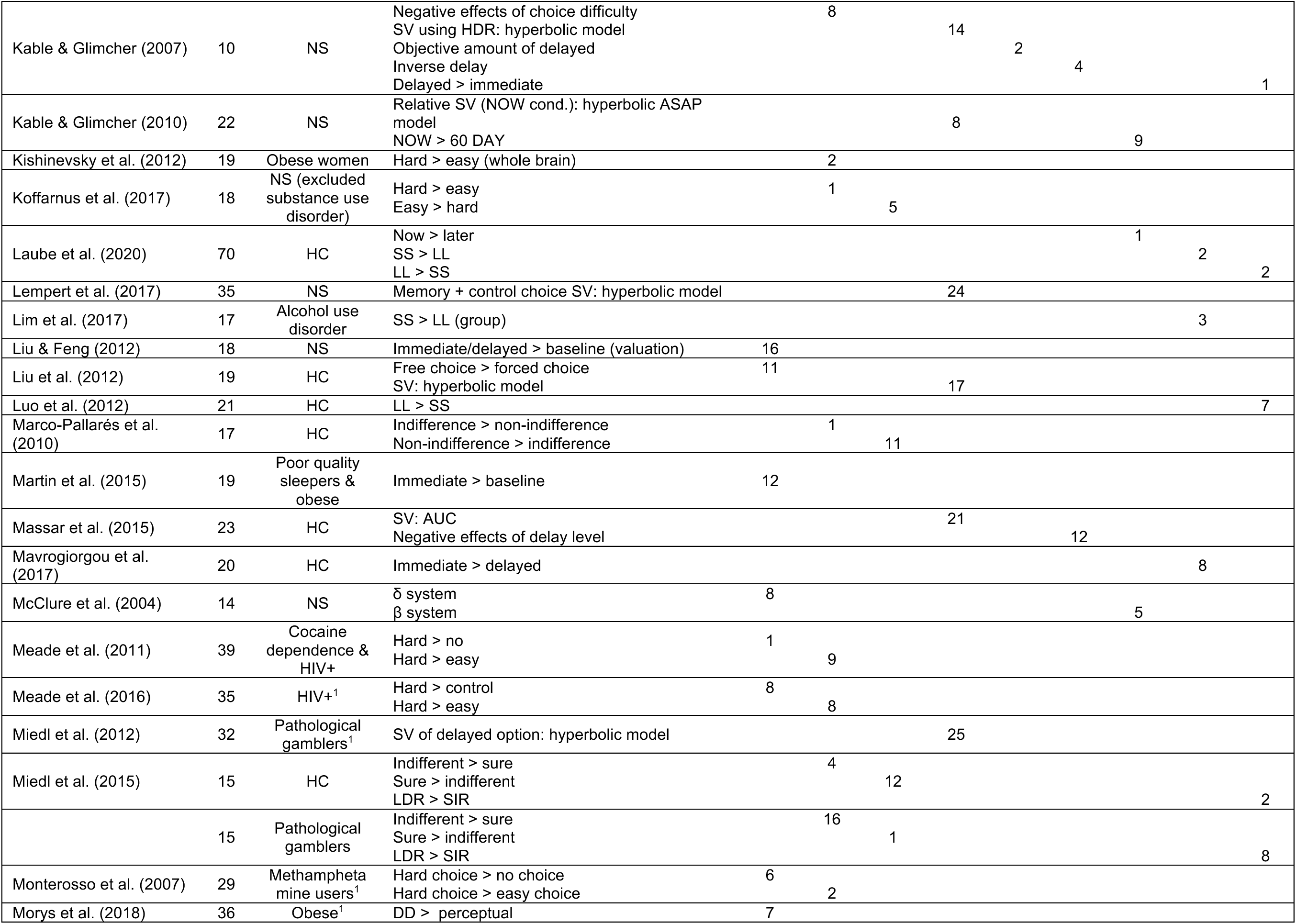
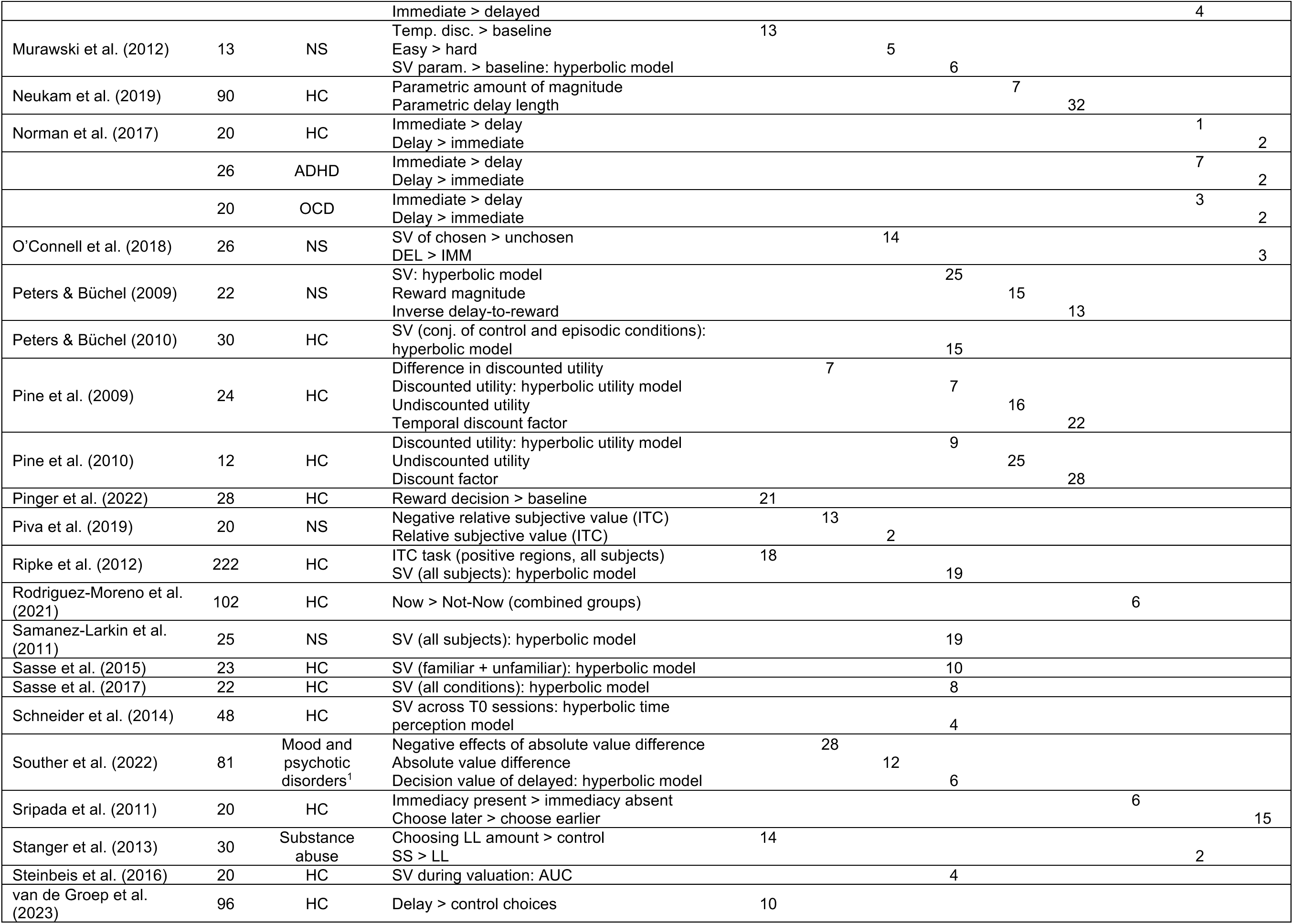
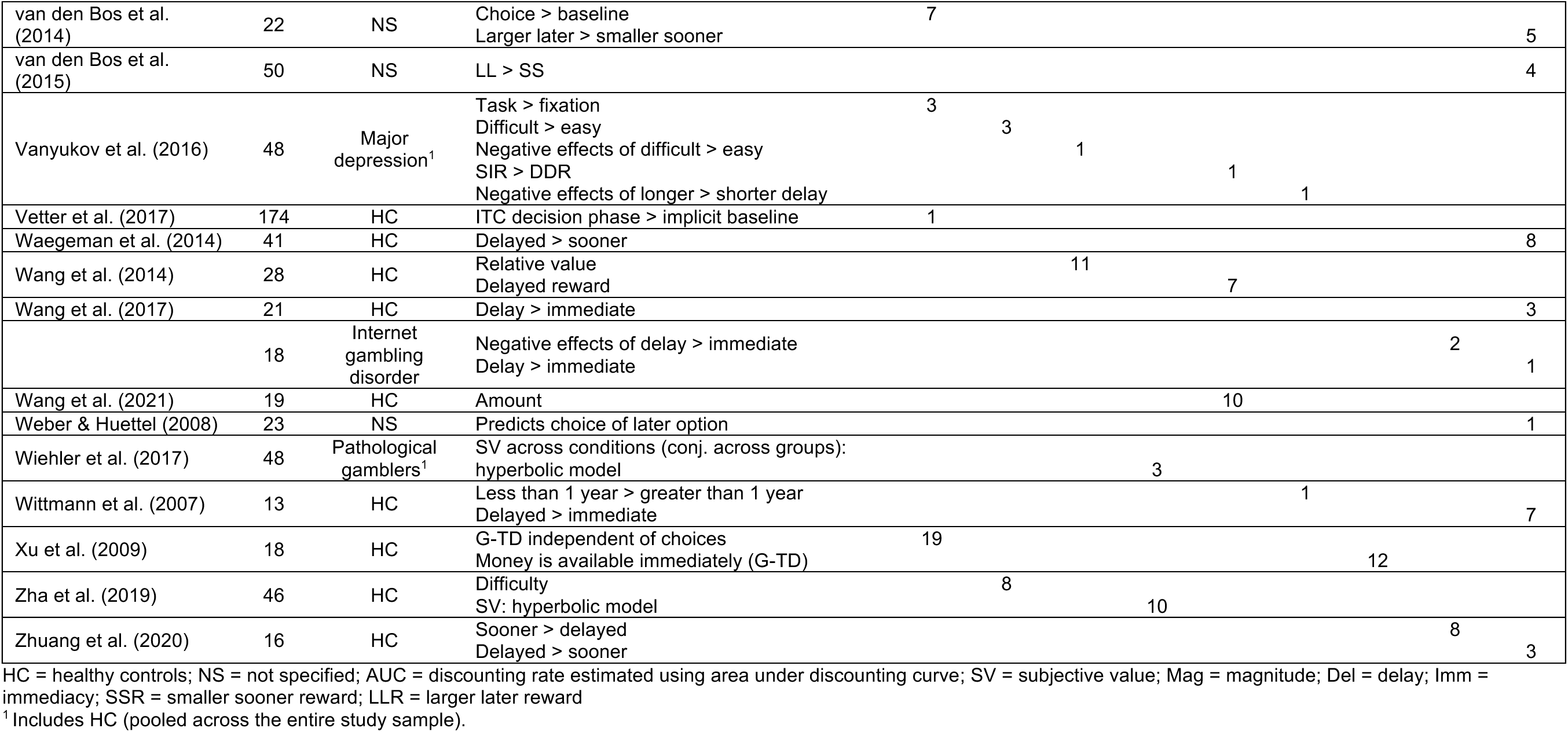
Meta-analytic sample (N = 80 studies; 150 contrasts).

### 2.3. Coding contrasts

Following study selection, we coded the studies into the following contrast categories: general task-related activity (*Task*), difficult decisions (*Hard*), easy decisions (*Easy*), subjective value (*SV*), objective amount (*Magnitude*), objective delay (*Delay*), choices involving an immediate reward (*Immediacy*), choosing the smaller-sooner reward (*SSR*), and choosing the larger-later reward (*LLR*).

For *Task*, we included positive effects of comparing the delay discounting task to a fixation or implicit baseline or a control task (e.g., choose left/right, choose larger). We also categorized as *Task* comparing a subset of the task to baseline (e.g., *Hard* > baseline). In cases where more than one subset was available (e.g., *Hard* > baseline and *Easy* > baseline), we included the contrast that would theoretically elicit a larger effect (i.e., *Hard* > baseline). Trials in which the two options are closer in subjective value or near the indifference point are considered to be difficult, and studies categorize trials as hard or easy based on the value distance or distance from the indifference point. Therefore, *Hard* included direct comparisons between hard and easy trials (positive effects of *Hard* > *Easy* or negative effects of *Easy* > *Hard*) whereas *Easy* included the opposite effects. We also coded negative correlations with unsigned difference in subjective value between the two options as *Hard* and positive correlations as *Easy*. For *SV*, we included brain regions showing positive correlation with the discounted value of the delayed reward. Although *SV* is frequently calculated using hyperbolic discounting (Commons et al., 2013), we deferred to the judgment of the original authors and reviewers and included all discounted value regressors regardless of how *SV* was calculated. For reference, we list the model used to calculate *SV* in each study in Table 1. For *Magnitude*, we included positive correlations with the objective amount of either the *SSR* or *LLR*, although all studies in our meta-analytic sample examined the amount of the delayed option except for one study (Albert, 2011), which examined varying amounts of the immediate option while keeping the amount of delayed reward constant throughout the task. For *Delay*, we included brain regions showing reduced activations as the delay increases (i.e., the negative effects of delay or positive effects of inverse delay). For *Immediacy*, we included comparing trials involving an immediate reward to either all trials or trials involving delayed options only. *SSR* included direct comparisons between the two choices (positive effects of *SSR* > *LLR* or negative effects of *LLR* > *SSR*) and *LLR* included the converse.

As previously noted, we did not exclude populations with psychiatric illnesses from our meta-analyses, and several of the studies in our sample report results separately for different psychiatric groups. Rather than selecting a single group from each study, we chose to include all groups for which a relevant contrast was reported, treating each as an independent study for the purpose of analysis. The alternative approach of merging foci from separate populations into one experiment would violate independence assumptions of MKDA’s random-effects model (Wager et al., 2009).

After coding, there were 29 studies (282 foci) in *Task*, 18 studies (140 foci) in *Hard*, 14 studies (107 foci) in *Easy*, 24 studies (275 foci) in *SV*, 10 studies (92 foci) in *Magnitude*, 9 studies (116 foci) in *Delay*, 9 studies (77 foci) in *Immediacy*, 14 studies (80 foci) in *SSR*, and 24 studies (142 foci) in *LLR*.

### 2.4. Coordinate-based meta-analysis

To synthesize results across studies and identify brain regions with consistent activation for each contrast, we used *coordinate-based meta-analysis* (CBMA). The neuroimaging findings we are synthesizing are reported as a set of stereotactic coordinates denoting the foci of peak activations for each contrast. CBMA focuses on the consistency of the spatial *location* of effects across the brain, unlike traditional meta-analyses in other disciplines that examine the magnitude of effect *size*. One such CBMA technique is *activation likelihood estimation* (ALE; Eickhoff et al., 2009, 2012; Turkeltaub et al., 2012), which was the tool of choice for some earlier meta-analyses of delay discounting (Carter et al., 2010; Wesley & Bickel, 2014). Given a set of studies, ALE uses a Gaussian kernel to model activation peaks as probability distributions, estimating the probability that at least one focus of activation truly lies in a given voxel. In the present meta-analysis, we primarily employ an alternative method used in a previous meta-analysis from our group (Bartra et al., 2013) based on multilevel kernel density analysis (MKDA; Wager et al. 2007). MKDA uses a uniform spherical kernel to calculate the proportion of studies with an activation focus lying near a given voxel. Both ALE and MKDA are considered suitable approaches to CBMA and have been shown to yield largely similar results (Eickhoff et al., 2012; Salimi-Khorshidi et al., 2009). Moreover, the modified MKDA procedure used here adapts the two-step cluster-level inference algorithm standard in ALE (Eickhoff et al., 2012). Despite these similarities, there are two advantages motivating the use of our method over ALE: (1) MKDA leads to a more interpretable test statistic—the percentage of studies reporting an effect within a pre-defined distance from a given voxel. (2) We used a probabilistic gray-matter mask for null-hypothesis testing (described in detail in section 2.5 below), instead of the more liberal binary gray-matter mask utilized in ALE (which retains 92.6% of the voxels of a whole-brain mask).

Nevertheless, ALE comes with its own advantages, including an empirically-derived kernel width based on study sample size (Eickhoff et al., 2009). As such, while MKDA is the primary analysis reported in this paper, we also conducted non-preregistered *post hoc* ALE analyses for all contrasts, which corroborate our main findings. These analyses were performed using GingerALE (v3.0.2; https://brainmap.org/ale/), setting an uncorrected threshold of *p* < 0.001 for 1000 null-hypothesis permutations, with a cluster-level threshold of *p* < 0.05.

Other techniques for CBMA include powerful automated pipelines such as Neurosynth (https://neurosynth.org/analyses/; Yarkoni et al., 2011). Neurosynth allows users to examine meta-analytic maps constructed from coordinates that appear in papers that contain terms of interest, such as “delay” or “value.” However, Neurosynth maps do not address the questions in the current study, which concern the consistency of activation observed for specific contrasts. To address these questions, manual curation is still required, especially as contrasts can be referred to by different names across studies.

### 2.5. Multilevel kernel density analysis

Taking each study’s coordinates of peak activation as input, MKDA quantifies the proportion of studies that report activation foci within a given radius (Wager et al., 2007). The specific MKDA procedure applied here was used in Bartra et al. (2013) and incorporates several adjustments to methods developed in Wager et al. (2007).

We first ensured all coordinates were in MNI space, converting those originally reported in Talairach space using the Lancaster transform (Lancaster et al., 2007) implemented in GingerALE (*tal2icbm_other*). For each study, we then created a binary indicator map (IM), assigning a value of 100 for voxels within 10mm of the foci and 0 for others. We averaged these maps across studies, resulting in the mean IM ranging from 0-100 that can be interpreted as the percentage of studies reporting a peak within a 10mm radius. For significance testing, we implemented permutation testing in MATLAB by simulating a null hypothesis that foci are randomly distributed, while maintaining the number of studies and number of foci per study. We also applied a cluster-forming threshold for each contrast to improve sensitivity. For each study, we first accounted for the probability of gray matter, where there is a higher probability of observing a positive IM value, by calculating the proportion of voxels with a positive IM value in the top quintile, as indicated by the ICBM tissue probability map (Mazziotta et al., 1995). These proportions were then treated as independent probabilities to derive the mean IM value that would be exceeded with a probability of less than 0.01 by chance alone. We applied this threshold to the results of each random permutation and recorded the largest cluster mass (sum of all IM values in a given cluster) in the null distribution. We also applied the same threshold to the real data, and the resulting clusters were assigned corrected p-values based on the frequency of equal or larger clusters in the null distribution. All reported results are based on 5000 permutation iterations (one of which was the observed data) and a threshold of corrected *p* = 0.05 (one-tailed).

## 3. Results

We first examined contrasts expected to identify activity in the frontoparietal and salience networks. We found significant clusters for both *Task* and *Hard* contrasts in these regions. For general task-related activations (*Task*), we found significant clusters in bilateral dorsomedial prefrontal cortex (dmPFC), bilateral insula, left middle frontal gyrus (MFG), and right frontal pole. We also observed clusters in bilateral occipital cortex and bilateral occipital poles. For choice difficulty (*Hard*), significant clusters were found in bilateral dmPFC and bilateral insula. For negative effects of difficulty (*Easy*), we found significant clusters in the left supramarginal gyrus and right PCC.

We then examined contrasts expected to identify activity in the valuation network, including *SV*, *Magnitude*, *Delay,* and *Immediacy*. The highest meta-analytic statistics (i.e., percentage of studies reporting a peak within 10mm from a given voxel) were found bilaterally in the striatum for *SV* (62.5%). Additionally, we observed significant clusters for *SV* in bilateral ventromedial prefrontal cortex (vmPFC), left PCC, and left middle temporal gyrus. *Magnitude* foci were also clustered in the right striatum. For negative effects of *Delay*, significant clustering was found in the right frontal pole, bilateral anterior cingulate cortex (ACC), and right MFG. We also found significant clusters for *Immediacy* in the left vmPFC. Taken together, these results suggest that *SV*, *Magnitude*, and *Immediacy* contrasts produce reliable activations in overlapping regions, namely the striatum and vmPFC.

Lastly, we examined contrasts between impulsive and future-oriented choices. As previously described, the CNDS framework predicts greater activations in the valuation network when choosing the smaller-sooner reward (*SSR > LLR*) and greater frontoparietal and salience activations when choosing the larger-later reward (*LLR > SSR*). Contrary to these predictions, we only observed a small cluster in the left dmPFC for *SSR*. For *LLR*, we only found clusters in the left precentral gyrus and right occipital fusiform gyrus. Given the localization of the largest effect for *LLR* in motor cortex, we examined the studies that contributed to this cluster for motor confounds and found that majority of studies that contributed to this cluster instructed participants to choose the *LLR* using their right middle finger (and *SSR* with the right index finger; Christakou et al., 2011; Claus et al., 2011; Miedl et al., 2015; Norman et al., 2017; Waegeman et al., 2014). Based on these findings, we conclude that choice contrasts do not reliably identify the predicted neural networks across experiments.

See Table 2 and Figure 2 for results for all contrasts.

**Table 2.**
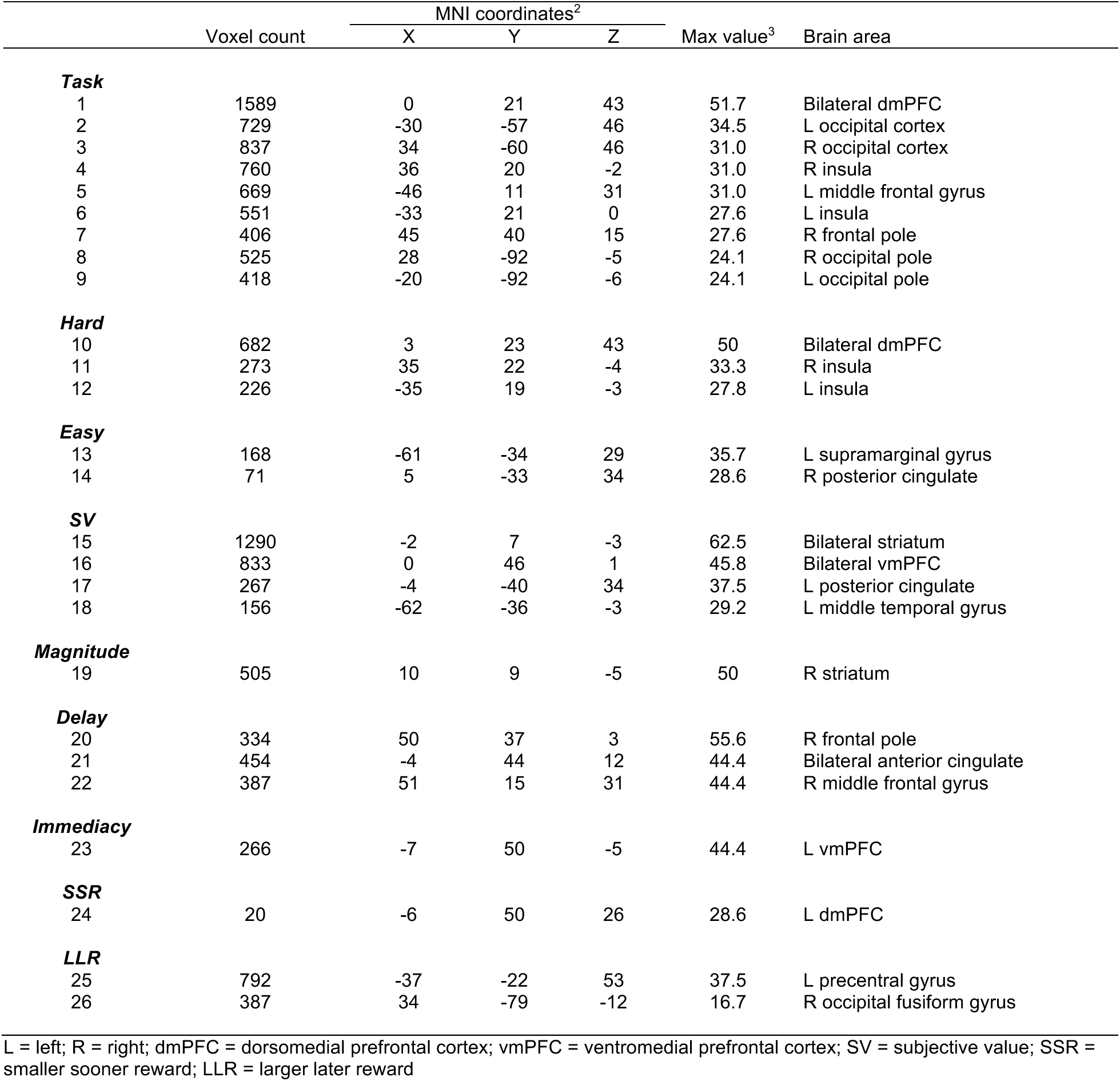
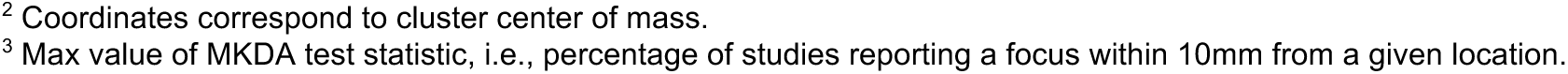
MKDA meta-analysis cluster list.

|  | Voxel count | MNI coordinates<sup>2</sup> |  |  | Max value<sup>3</sup> | Brain area |
| --- | --- | --- | --- | --- | --- | --- |
|  |  | X | Y | Z |  |  |
| **Task** |  |  |  |  |  |  |
| 1 | 1589 | 0 | 21 | 43 | 51.7 | Bilateral dmPFC |
| 2 | 729 | -30 | -57 | 46 | 34.5 | L occipital cortex |
| 3 | 837 | 34 | -60 | 46 | 31.0 | R occipital cortex |
| 4 | 760 | 36 | 20 | -2 | 31.0 | R insula |
| 5 | 669 | -46 | 11 | 31 | 31.0 | L middle frontal gyrus |
| 6 | 551 | -33 | 21 | 0 | 27.6 | L insula |
| 7 | 406 | 45 | 40 | 15 | 27.6 | R frontal pole |
| 8 | 525 | 28 | -92 | -5 | 24.1 | R occipital pole |
| 9 | 418 | -20 | -92 | -6 | 24.1 | L occipital pole |
| **Hard** |  |  |  |  |  |  |
| 10 | 682 | 3 | 23 | 43 | 50 | Bilateral dmPFC |
| 11 | 273 | 35 | 22 | -4 | 33.3 | R insula |
| 12 | 226 | -35 | 19 | -3 | 27.8 | L insula |
| **Easy** |  |  |  |  |  |  |
| 13 | 168 | -61 | -34 | 29 | 35.7 | L supramarginal gyrus |
| 14 | 71 | 5 | -33 | 34 | 28.6 | R posterior cingulate |
| **SV** |  |  |  |  |  |  |
| 15 | 1290 | -2 | 7 | -3 | 62.5 | Bilateral striatum |
| 16 | 833 | 0 | 46 | 1 | 45.8 | Bilateral vmPFC |
| 17 | 267 | -4 | -40 | 34 | 37.5 | L posterior cingulate |
| 18 | 156 | -62 | -36 | -3 | 29.2 | L middle temporal gyrus |
| **Magnitude** |  |  |  |  |  |  |
| 19 | 505 | 10 | 9 | -5 | 50 | R striatum |
| **Delay** |  |  |  |  |  |  |
| 20 | 334 | 50 | 37 | 3 | 55.6 | R frontal pole |
| 21 | 454 | -4 | 44 | 12 | 44.4 | Bilateral anterior cingulate |
| 22 | 387 | 51 | 15 | 31 | 44.4 | R middle frontal gyrus |
| **Immediacy** |  |  |  |  |  |  |
| 23 | 266 | -7 | 50 | -5 | 44.4 | L vmPFC |
| **SSR** |  |  |  |  |  |  |
| 24 | 20 | -6 | 50 | 26 | 28.6 | L dmPFC |
| **LLR** |  |  |  |  |  |  |
| 25 | 792 | -37 | -22 | 53 | 37.5 | L precentral gyrus |
| 26 | 387 | 34 | -79 | -12 | 16.7 | R occipital fusiform gyrus |

L = left; R = right; dmPFC = dorsomedial prefrontal cortex; vmPFC = ventromedial prefrontal cortex; SV = subjective value; SSR = smaller sooner reward; LLR = larger later reward
<sup>2</sup> Coordinates correspond to cluster center of mass.
<sup>3</sup> Max value of MKDA test statistic, i.e., percentage of studies reporting a focus within 10mm from a given location.

**Fig. 2.**
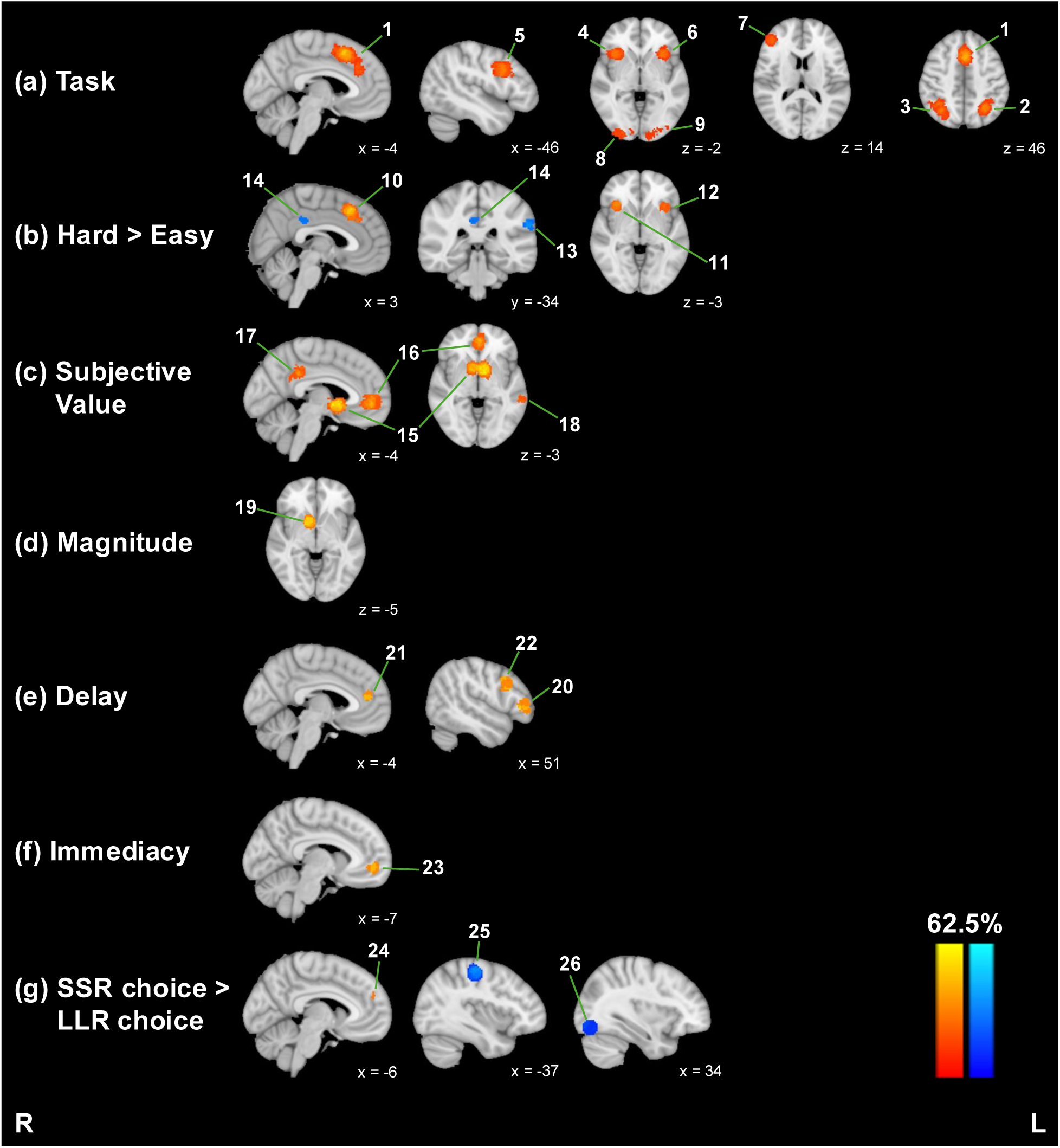
MKDA meta-analysis results. Warm colors represent positive effects, and cool colors represent negative effects. Numbered labels correspond to cluster numbers in Table 2.

### 3.1. Robustness check: meta-analytic method

To test whether the results above depended on the meta-analytic method we chose, we repeated meta-analysis of each contrast using ALE. With ALE, we see largely parallel results to those reported above, including for the *SSR* and *LLR* contrasts. See Supplementary Table 1 and Supplementary Figure 1 for a full list of significant clusters for all nine contrasts.

### 3.2. Robustness check: inclusion of clinical samples

We also tested whether the results above depended on our choice to include studies of populations with psychiatric disorders and other clinical diagnoses (24.7% of studies included). It is possible that the inclusion of clinical studies increased heterogeneity and reduced the consistency observed. We re-performed the meta-analyses above, excluding all studies of clinical populations (i.e., any group with a label other than *HC* or *NS* under the *Diagnosis* column in Table 1). We performed both MKDA and ALE. Once again, we find qualitatively similar results across the board, including a lack of reliable activations in predicted regions for the choice contrasts. See Supplementary Tables 2 and 3 and Supplementary Figures 2 and 3 for a full report.

## 4. DISCUSSION

In this paper, we report a series of meta-analyses examining neural effects revealed by commonly used analytic approaches in fMRI studies of delay discounting. Specifically, we investigated the consistency of neural activations in the valuation and frontoparietal and salience networks. Despite its wide use, our results did not reveal reliable effects of choice (*SSR* or *LLR*) in either of these two systems. Instead, the most reliable clusters were found throughout the valuation network as a function of subjective value. Although the objective amount and presence of an immediate option engaged similar regions such as the striatum and vmPFC, we saw more widespread activations for subjective value than the other contrasts. In addition, the frontoparietal and salience networks were reliably engaged by task and choice difficulty contrasts. Taken together, these results offer meta-analytic support for examining value-based contrasts rather than comparing choices. In other words, the hypothesis that there are distinct neural systems associated with impulsive or patient choices is not supported by our work.

This conclusion is further bolstered by considering the number of studies that reported null findings. Because MKDA estimates the location of effects rather than the size of effects, null findings are not included and there is no statistical penalty for null findings (Fox et al., 1998; Wager et al., 2007). Given this, it is worth noting that, during our study search, we identified 20 studies reporting no significant clusters for the *SSR > LLR* contrast (Fu et al., 2022; Hamilton et al., 2020; Hare et al., 2014; Kable & Glimcher, 2007; Koffarnus et al., 2017; Luo et al., 2012; Marco-Pallarés et al., 2010; Martin et al., 2015; Miedl et al., 2015; Monterosso et al., 2007; O’Connell et al., 2018; Pinger et el., 2022; Schmaal et al., 2014; Sripada et al., 2011; van de Groep et al., 2023; van den Bos et al., 2014, 2015; Waegeman et al., 2014; Weber & Huettel, 2008; Wittmann et al., 2007) and 15 studies with no significant clusters for the *LLR > SSR* contrast (Albrecht et al., 2013; Hamilton et al., 2020; Kable & Glimcher, 2010; Kobiella et al., 2014; Koffarnus et al., 2017; Lim et al., 2017; Marco-Pallarés et al., 2010; Martin et al., 2015; Mavrogiorgou et al., 2017; Monterosso et al., 2007; Morys et al., 2018; Pinger et al., 2022; Schmaal et al., 2014; Stanger et al., 2013; van de Groep et al., 2023). In comparison, only two studies reported null findings for SV (Miedl et al., 2014; Sripada et al., 2011).

### 4.1. Comparison to other meta-analyses

Our results are broadly in line with early meta-analytic work which identified widespread regions in medial and lateral prefrontal cortex, PCC, insula, and striatum as being engaged during delay discounting (Carter et al., 2010; Wesley & Bickel, 2014). However, these foundational reviews did not separately examine activity associated with different contrasts. Our findings demonstrate how these different regions are engaged during delay discounting, with the valuation network (striatum, vmPFC, PCC) being engaged as a function of the subjective value of the rewards, and the frontoparietal (lateral prefrontal and lateral parietal) and salience (ACC and insula) networks being engaged more during delay discounting than baseline or control tasks, and increasingly so with choice difficulty. Since we began this work, a similar meta-analysis comparing a subset of these contrasts in healthy adults has been published (Schüller et al., 2019). In that study, the sample sizes for each contrast ranged from 3 to 13 experiments, which is far below current recommendations for meta-analytic sample size (Eickhoff et al., 2016). We had a much larger meta-analytic sample (9 to 29 experiments per contrast). Our results were nevertheless largely consistent with Schüller et al. (2019) for *Task*, *Hard*, *SV*, *Magnitude*, and *Immediacy* contrasts. However, contradicting Schüller et al., we did not find the expected distinct engagement of brain regions when making impulsive or future-oriented choices (*SSR* or *LLR*), despite including more than three times the sample size for *LLR* (n = 7 in Schüller et al., n = 24 in the present study). It is important to note that Schüller et al. used ALE and did not include clinical studies. However, we performed *post hoc* robustness checks and found that our main results were not dependent on either of these choices. We thus conclude the contradictory findings between our meta-analysis and Schüller et al. cannot be solely attributed to methodological differences and/or inclusion of clinical populations. Instead, we credit nearly five more years of new literature, as well as a generally more exhaustive sample size.

### 4.2. Clinical considerations

Despite our finding that choice-based contrasts do not reliably produce the theoretically predicted effects, this approach remains one of the most widely used in the literature. This is especially pronounced in clinical neuroimaging studies of delay discounting. Indeed, a large number of studies included in our meta-analysis used delay discounting to investigate a wide range of psychiatric disorders and related symptoms, such as substance use (Claus et al., 2011; Halcomb et al., 2022; Hoffman et al., 2008; Lim et al., 2017; Meade et al., 2011; Monterosso et al., 2007; Stanger et al., 2013), pathological gambling (Miedl et al., 2012, 2015; Wang et al., 2017; Wiehler et al., 2017), obsessive compulsive disorder (Norman et al., 2017), ADHD (Norman et al., 2017), schizophrenia (Avsar et al., 2013; Souther et al., 2022), major depressive disorder (Souther et al., 2022; Vanyukov et al., 2016), bipolar disorder (Souther et al., 2022), obesity (Kishinevsky et al., 2012; Martin et al., 2015; Morys et al., 2018), prediabetes (Deshpande et al., 2019), insomnia (Martin et al., 2015), and HIV (Meade et al., 2011, 2016).

Among the studies with a clinical sample included in our analyses, *SSR* (n = 7) was the third most frequently used contrast, whereas only three studies (Miedl et al., 2012; Souther et al., 2022; Wiehler et al., 2017) examined the *SV* contrast, and only one study (Vanyukov et al., 2016) examined the effects of *Magnitude* and *Delay*. This suggests that many existing clinical studies (and about half of basic decision neuroscience studies) have used an analytic strategy that has poor reliability. Therefore, we strongly recommend that future neuroimaging studies of delay discounting use reliable contrasts such as subjective value (to identify the valuation network) and choice difficulty (to identify the frontoparietal and salience networks).

### 4.3. Limitations

There are several limitations to consider when interpreting our findings.

First, although the contrasts related to our key findings (e.g., *LLR*, *SV*) had large sample sizes, some contrasts (e.g., *Magnitude*, *Delay*, *Immediacy*) had relatively smaller sample sizes. Future work should continue to investigate these understudied contrasts in order to identify analytic strategies that reliably reveal neural activity during delay discounting.

Second, for studies in our analysis reporting coordinate maps for independent groups of subjects (e.g., a clinical group and healthy controls), we chose to include *both* groups, treating each as a separate study, rather than selecting a single set of results to be included. Because these groups come from the same experiment and are not wholly independent, it is possible such studies may have had undue influence on our results. However, our *post hoc* analysis excluding clinical populations helps mitigate this concern. Likewise, most studies contributed more than one contrast, and these contrasts tended to go hand in hand (e.g., *SSR* and *LLR*, *Magnitude* and *Delay*, *Hard* and *Easy*). Similarly, because authors tended to use the same contrasts across multiple studies, there were several studies from the same research group in a given contrast.

Moreover, our study exclusively focused on delay discounting of monetary rewards. A limited but growing body of work has investigated neural activity underlying discounting of non-monetary rewards, revealing similar activation of the valuation, frontoparietal, and salience regions across reward types, including juice/water (McClure et al., 2007), arousing images (Prévost et al., 2010), and cigarettes (MacKillop et al., 2012). This dovetails with meta-analytic evidence supporting a common neural valuation system across commodities (Bartra et al., 2013; Levy & Glimcher, 2012), albeit with some subregional variation (Clithero & Rangel, 2014). We thus remain cautiously optimistic that our results are generalizable to other reward types. Nevertheless, at least one study of non-monetary delay discounting reports significant activations for choice-based contrasts (MacKillop et al., 2012), and several have found additional reward-specific activations (MacKillop et al., 2012; Markman et al., 2024), underscoring the need for more empirical work in this domain.

### 4.4. Future directions

The present meta-analysis offers several exciting directions for future research.

As data archiving becomes more common in the field, future meta-analyses can move beyond the coordinate-based meta-analysis approach and conduct image-based meta-analyses (IMBA; Salimi-Khorshidi et al., 2009). Unlike coordinate-based methods, which only include peak voxels, IMBA includes the whole statistical map, allowing researchers to detect consistent but relatively weaker effects and to calculate effect sizes, which is a limitation of coordinate-based methods that only examine effect locations. While the ideal meta-analysis would consist entirely of whole-brain statistical maps for a large sample of component studies, a more realistic hybrid approach known as effect-size seed-based *d* mapping (ES-SDM; Radua et al., 2012) incorporates peak coordinates as well as t-maps, when available. We encourage researchers to make whole-brain statistical maps publicly available in established archives, such as NeuroVault (https://neurovault.org/).

Future meta-analytic work in delay discounting might also aim to examine the link between these neural effects and individual differences, incorporating behavioral indicators such as discount rate and/or cognitive measures. Similarly, as more neuroimaging studies of delay discounting in clinical populations become available, it would be worthwhile to conduct a future meta-analysis, both within specific disorders and across psychopathology in line with the RDoC framework. Future meta-analyses could also expand on our work and include paradigms that are tailored to clinical relevance (e.g., cigarettes for smokers; MacKillop et al., 2012; food for obese individuals; Weygandt et al., 2015, 2019).

Additionally, it would be important for future research to focus on how the valuation, frontoparietal, and salience networks jointly shape discounting. Although our results are not compatible with the CNDS framework, all three networks were still reliably engaged by one or another contrast. Recruitment of frontoparietal and salience networks during *Task* and *Hard* contrasts is consistent with domain-general functions, such as conflict monitoring and cognitive control (Botvinick et al., 2001). Significant activation in dmPFC (and nearby dorsal ACC) for the *Hard* contrast also fits nicely with more recent theoretical work implicating this region in the allocation of cognitive resources for effortful tasks (Shenhav et al., 2016). Intriguingly, others have offered a special role for these networks in decision-making, such as value comparison (Kable & Glimcher, 2009) and value accumulation (Rodriguez et al., 2015). Examining functional connectivity patterns during delay discounting could help tease apart these possibilities. An increasing number of studies have taken this approach (e.g., Hare et al., 2014; van den Bos et al., 2014), offering another promising direction for future meta-analyses.

A practical goal of our work was to identify analytic strategies that are reliable and robust and provide recommendations for future research. Given the high reliability and interpretability of effects produced by the subjective value and choice difficulty contrasts, our recommendation for future work is to use these analytic approaches in the absence of specific hypotheses that call for a different approach. Another practical outcome of our work is the generation of cluster maps that can be used as *a priori* regions of interest for future experiments. This would be especially valuable for clinical neuroimaging studies of delay discounting that might be underpowered to detect modest effects using whole-brain approaches. All results will be made publicly available for this purpose.

## 5. CONCLUSION

We found evidence of consistent neural activations in the valuation and frontoparietal and salience networks during delay discounting. The valuation network was most reliably activated by subjective value and was also reliably engaged by objective magnitude and reward immediacy. The frontoparietal and salience networks were reliably activated by task and choice difficulty contrasts. Despite its wide use, our results did not reveal reliable effects of choosing the larger later or smaller sooner options in either of these two systems. Therefore, we strongly recommend that future neuroimaging studies of delay discounting use analytic approaches shown to reliably identify specific networks (e.g., subjective value for the valuation network, choice difficulty for the frontoparietal and salience networks).

## DATA AND CODE AVAILABILITY

All coordinate files, code, results, and cluster maps are available on the OSF: https://osf.io/xvt5k/.

## AUTHOR CONTRIBUTIONS

N.V.L.: Data curation, Formal analysis, Investigation, Methodology, Visualization, Writing – review & editing. M.K.S.: Conceptualization, Data curation, Formal analysis, Investigation, Methodology, Writing – original draft, Writing – review & editing. B.B.: Data curation, Formal analysis, Investigation, Writing – review & editing. J.W.K.: Conceptualization, Funding acquisition, Investigation, Methodology, Resources, Supervision, Writing – original draft, Writing – review & editing.

## SUPPLEMENTARY MATERIALS

A document containing supplementary analyses is provided along with this pre-print.

