## Supplementary Material for "A meta-analysis of neural systems underlying delay discounting: Implications for transdiagnostic research"

**Supplementary Table 1.** GingerALE meta-analysis cluster list.

| | | MNI coordinates[1] | | | | |
|---|---|---|---|---|---|---|
| | Voxel count | X | Y | Z | Max value[2] | Brain area |
| ***Task*** | | | | | | |
| 1 | 714 | 0 | 21 | 44 | 0.045 | Bilateral dmPFC |
| 2 | 274 | 37 | 20 | -2 | 0.0376 | R insula |
| 3 | 270 | -46 | 10 | 31 | 0.0357 | L middle frontal gyrus |
| 4 | 245 | -33 | 21 | 0 | 0.0316 | L insula |
| 5 | 152 | 46 | 40 | 15 | 0.031 | R frontal pole |
| 6 | 308 | -29 | -57 | 47 | 0.029 | L occipital cortex |
| 7 | 123 | -16 | -93 | -8 | 0.0269 | L occipital pole |
| 8 | 290 | 33 | -61 | 46 | 0.0263 | R occipital cortex |
| ***Hard*** | | | | | | |
| 9 | 455 | 3 | 24 | 43 | 0.0259 | Bilateral dmPFC |
| 10 | 175 | -34 | 19 | -2 | 0.0227 | L insula |
| 11 | 184 | 35 | 22 | -4 | 0.0206 | R insula |
| 12 | 88 | 1 | 0 | 29 | 0.018 | Bilateral anterior cingulate |
| ***Easy*** | | | | | | |
| 13 | 143 | -62 | -33 | 28 | 0.0212 | L supramarginal gyrus |
| 14 | 69 | -58 | -7 | -10 | 0.0164 | L superior temporal gyrus |
| 15 | 85 | 5 | -32 | 35 | 0.0143 | R posterior cingulate |
| ***SV*** | | | | | | |
| 16 | 787 | -2 | 8 | -3 | 0.0636 | Bilateral striatum |
| 17 | 479 | 1 | 47 | 1 | 0.0346 | Bilateral vmPFC |
| 18 | 183 | -63 | -35 | -4 | 0.0274 | L middle temporal gyrus |
| 19 | 185 | -4 | -40 | 34 | 0.0202 | L posterior cingulate |
| ***Magnitude*** | | | | | | |
| 20 | 180 | 10 | 8 | -5 | 0.0265 | R striatum |
| ***Delay*** | | | | | | |
| 21 | 91 | 52 | 16 | 31 | 0.0193 | R middle frontal gyrus |
| 22 | 131 | 49 | 37 | 2 | 0.017 | R frontal pole |
| 23 | 98 | -9 | 44 | 15 | 0.0152 | L anterior cingulate |
| ***Immediacy*** | | | | | | |
| 24 | 136 | -6 | 52 | -6 | 0.0172 | L vmPFC |
| ***SSR*** | | | | | | |
| 25 | 62 | -4 | 55 | 25 | 0.0135 | L dmPFC |
| ***LLR*** | | | | | | |
| 26 | 268 | -37 | -22 | 53 | 0.0344 | L precentral gyrus |
| 27 | 87 | 34 | -78 | -12 | 0.0238 | R occipital fusiform gyrus |
| 28 | 100 | -23 | -85 | -10 | 0.0218 | L occipital fusiform gyrus |

L = left; R = right; dmPFC = dorsomedial prefrontal cortex; vmPFC = ventromedial prefrontal cortex; SV = subjective value; SSR = smaller sooner reward; LLR = larger later reward
[1] Coordinates correspond to cluster center of mass.
[2] Max value of ALE test statistic, i.e., probability that at least one focus of activation truly lies in a given location.

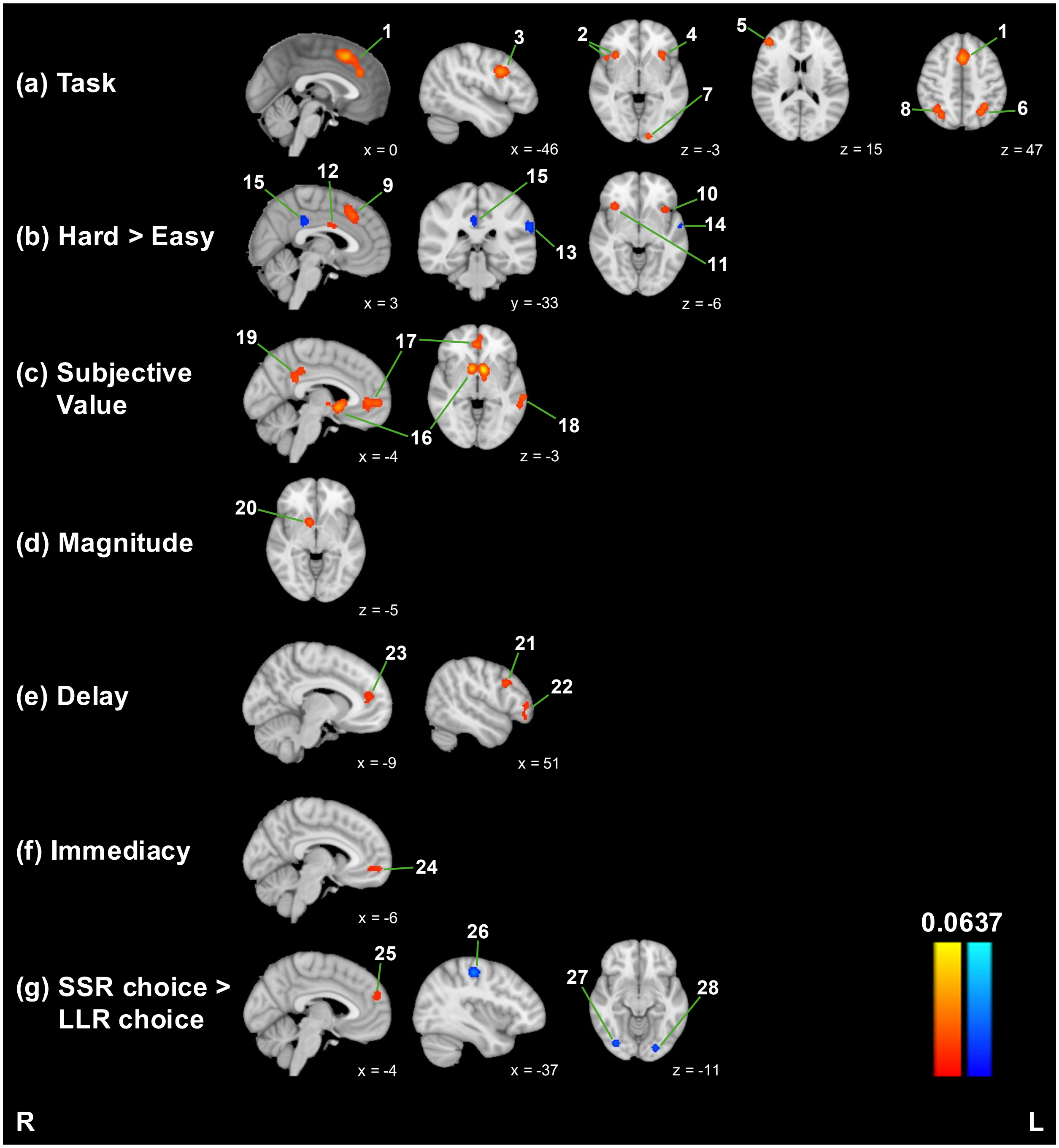


L = left; R = right; SSR = smaller sooner reward; LLR = larger later reward

**Supplementary Fig. 1.** GingerALE meta-analysis results. Warm colors represent positive effects, and cool colors represent negative effects. Numbered labels correspond to cluster numbers in Supplementary Table 1.

**Supplementary Table 2.** MKDA meta-analysis cluster list, excluding studies with clinical populations.

| | Volume ($mm^3$) | MNI coordinates[3] X | Y | Z | Max value[4] | Brain area |
|---|---|---|---|---|---|---|
| ***Task*** | | | | | | |
| 1 | 688 | -2 | 15 | 50 | 57.9 | Bilateral dmPFC |
| 2 | 542 | -29 | -58 | 46 | 42.1 | L occipital cortex |
| 3 | 382 | 34 | -59 | 45 | 36.8 | R occipital cortex |
| 4 | 344 | -45 | 11 | 30 | 31.6 | L middle frontal gyrus |
| 5 | 334 | -33 | 21 | 0 | 31.6 | L insula |
| 6 | 230 | 34 | 20 | 0 | 31.6 | R insula |
| 7 | 223 | 46 | 41 | 15 | 31.6 | R frontal pole |
| ***Hard*** | | | | | | |
| 8 | 930 | 4 | 24 | 44 | 50 | Bilateral dmPFC |
| ***Easy*** | | | | | | |
| | | | No significant clusters | | | |
| ***SV*** | | | | | | |
| 9 | 1472 | -3 | 6 | -3 | 61.9 | Bilateral striatum |
| 10 | 1067 | 2 | 45 | 3 | 38.1 | Bilateral vmPFC |
| 11 | 312 | -5 | -40 | 35 | 33.3 | L posterior cingulate |
| ***Magnitude*** | | | | | | |
| 12 | 505 | 10 | 9 | -5 | 55.6 | R striatum |
| ***Delay*** | | | | | | |
| 13 | 334 | 50 | 37 | 3 | 62.5 | R frontal pole |
| 14 | 454 | -4 | 44 | 12 | 50 | Bilateral anterior cingulate |
| 15 | 387 | 51 | 15 | 31 | 50 | R middle frontal gyrus |
| ***Immediacy*** | | | | | | |
| 16 | 1554 | -4 | 49 | 2 | 50 | Bilateral vmPFC / anterior cingulate |
| 17 | 588 | 2 | 9 | -3 | 37.5 | Bilateral striatum |
| ***SSR*** | | | | | | |
| 18 | 444 | -5 | 55 | 24 | 42.9 | Bilateral dmPFC |
| ***LLR*** | | | | | | |
| 19 | 208 | -36 | -21 | 54 | 31.6 | L precentral gyrus |

L = left; R = right; dmPFC = dorsomedial prefrontal cortex; vmPFC = ventromedial prefrontal cortex; SV = subjective value; SSR = smaller sooner reward; LLR = larger later reward

[3] Coordinates correspond to cluster center of mass.

[4] Max value of MKDA test statistic, i.e., percentage of studies reporting a focus within 10mm from a given location.

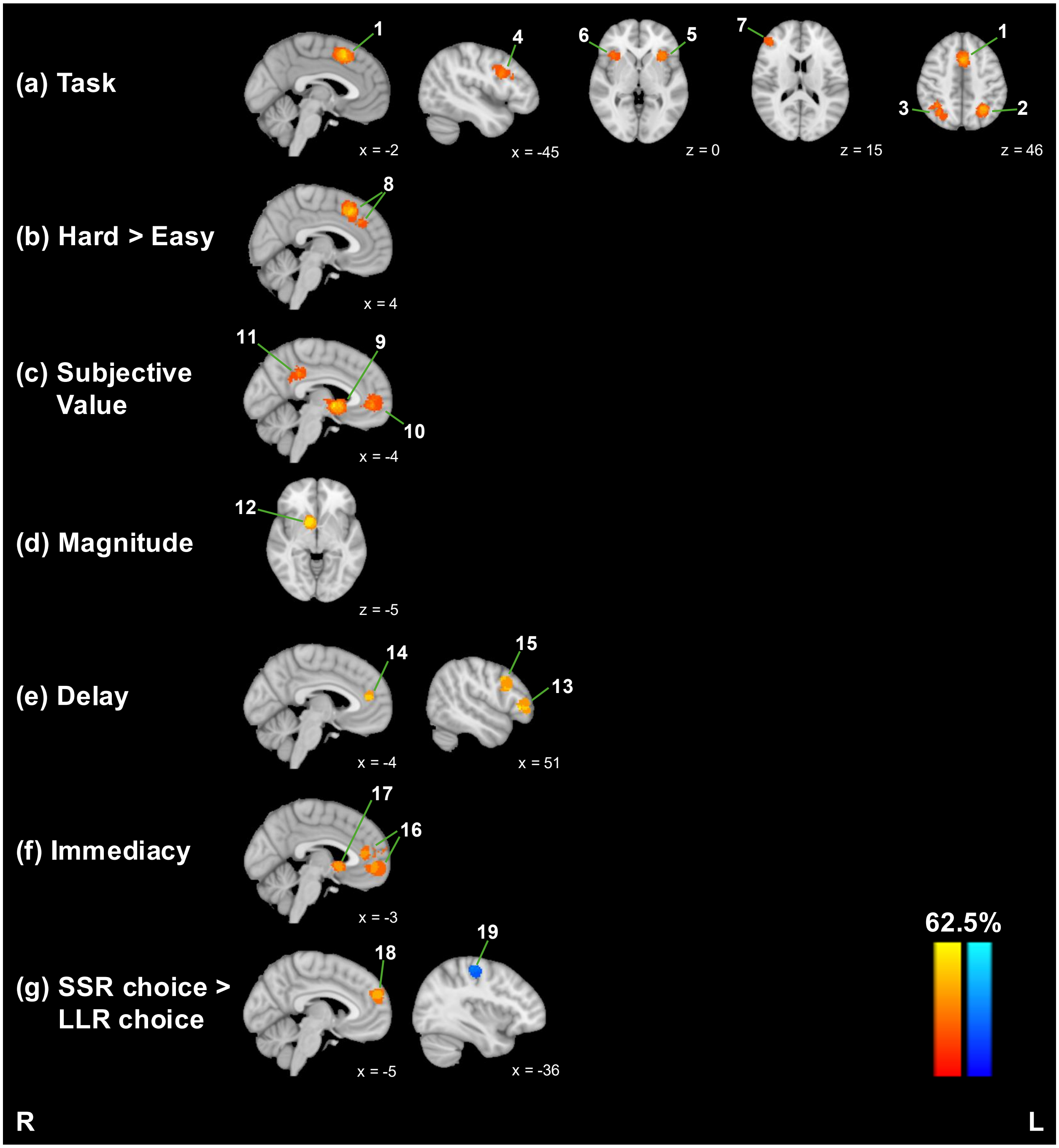


L = left; R = right; SSR = smaller sooner reward; LLR = larger later reward

**Supplementary Fig. 2.** MKDA meta-analysis results, excluding studies with clinical populations. Warm colors represent positive effects, and cool colors represent negative effects. Numbered labels correspond to cluster numbers in Supplementary Table 2.

**Supplementary Table 3.** GingerALE meta-analysis cluster list, excluding studies with clinical populations.

| | Voxel count | MNI coordinates[5] X | Y | Z | Max value[6] | Brain area |
|---|---|---|---|---|---|---|
| ***Task*** | | | | | | |
| 1 | 373 | -2 | 15 | 50 | 0.0317 | Bilateral dmPFC |
| 2 | 287 | -29 | -58 | 47 | 0.028 | L occipital cortex |
| 3 | 206 | -33 | 21 | 0 | 0.0246 | L insula |
| 4 | 110 | 34 | 20 | 1 | 0.023 | R insula |
| 5 | 97 | 47 | 41 | 16 | 0.0217 | R frontal pole |
| 6 | 141 | -45 | 10 | 30 | 0.0201 | L middle frontal gyrus |
| 7 | 151 | 33 | -62 | 45 | 0.0188 | R occipital cortex |
| ***Hard*** | | | | | | |
| 8 | 169 | 5 | 21 | 47 | 0.0179 | R dmPFC |
| 9 | 133 | 34 | 23 | -7 | 0.0166 | R insula |
| 10 | 113 | 44 | -41 | 44 | 0.0151 | R supramarginal gyrus |
| ***Easy*** | | | | | | |
| 11 | 87 | 4 | 15 | -7 | 0.0154 | R striatum |
| 12 | 92 | -62 | -33 | 29 | 0.013 | L supramarginal gyrus |
| 13 | 76 | 4 | -32 | 32 | 0.0125 | R posterior cingulate |
| ***SV*** | | | | | | |
| 14 | 689 | -2 | 7 | -3 | 0.0497 | Bilateral striatum |
| 15 | 324 | 3 | 46 | 1 | 0.0226 | Bilateral vmPFC |
| 16 | 109 | -54 | -58 | 20 | 0.0205 | L angular gyrus |
| 17 | 100 | -5 | -41 | 35 | 0.0168 | L posterior cingulate |
| ***Magnitude*** | | | | | | |
| 18 | 181 | 10 | 8 | -5 | 0.0265 | R striatum |
| ***Delay*** | | | | | | |
| 19 | 91 | 52 | 16 | 31 | 0.0193 | R middle frontal gyrus |
| 20 | 131 | 49 | 37 | 2 | 0.017 | R frontal pole |
| 21 | 98 | -9 | 44 | 15 | 0.0152 | L anterior cingulate |
| 22 | 64 | -9 | 12 | -1 | 0.0143 | L striatum |
| ***Immediacy*** | | | | | | |
| 23 | 160 | -6 | 52 | -6 | 0.0172 | L vmPFC |
| ***SSR*** | | | | | | |
| 24 | 98 | -4 | 55 | 25 | 0.0134 | L dmPFC |
| ***LLR*** | | | | | | |
| 25 | 157 | -35 | -22 | 53 | 0.0204 | L precentral gyrus |

L = left; R = right; dmPFC = dorsomedial prefrontal cortex; vmPFC = ventromedial prefrontal cortex; SV = subjective value; SSR = smaller sooner reward; LLR = larger later reward

[5] Coordinates correspond to cluster center of mass.

[6] Max value of ALE test statistic, i.e., probability that at least one focus of activation truly lies in a given location.

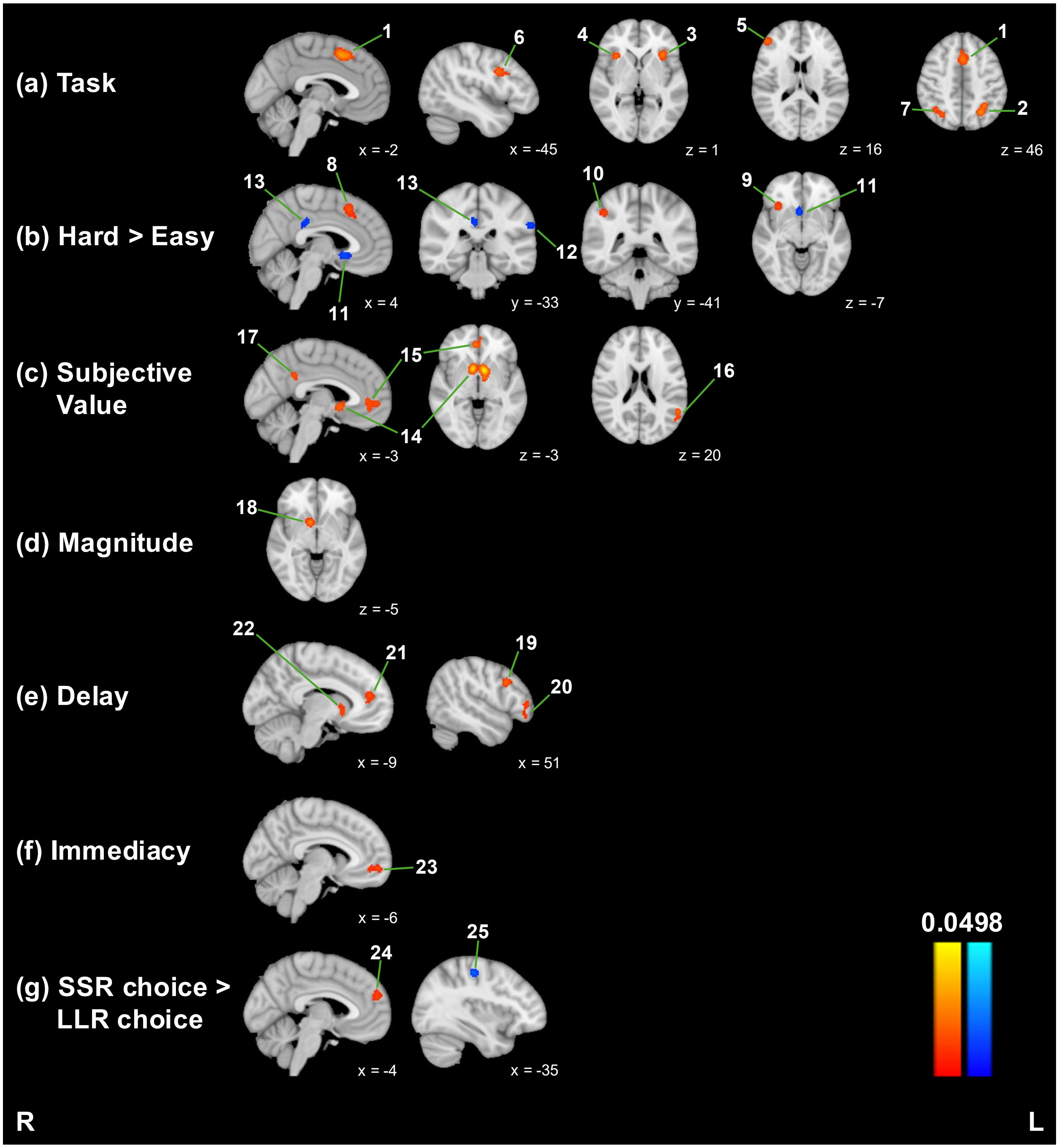


L = left; R = right; SSR = smaller sooner reward; LLR = larger later reward

**Supplementary Fig. 3.** GingerALE meta-analysis results, excluding studies with clinical populations. Warm colors represent positive effects, and cool colors represent negative effects. Numbered labels correspond to cluster numbers in Supplementary Table 3.
